# A double-spiral maze and hi-resolution tracking pipeline to study dispersal by groups of minute insects

**DOI:** 10.1101/2022.09.06.506709

**Authors:** M. Cointe, V. Burte, G. Perez, L. Mailleret, V. Calcagno

## Abstract

Minute insects such as parasitic micro-wasps have high basic and applied importance, for their widespread use as biocontrol agents. Their dispersal is a phenotype of particular interest. Classically, it is evaluated using field releases, but those are time consuming, costly, and their results highly variable, preventing high-throughput and repeatability. Alternatively, dispersal can be studied using small-scale assays, but those neglect important higher-scale processes. Consequently, proper evaluation of dispersal is often complicated or lacking in academic studies and biocontrol breeding programs. Here we introduce a new method, the double-spiral maze, that allows the study of spatial propagation at relevant scales (several hours and meters), retaining high throughput and experimental power. The method records the location of every individual at every time, enabling accurate precise estimates of diffusion coefficients or other dispersal metrics. We describe this affordable, scalable, and easy-to-implement method, and illustrate its application with a species of agricultural interest.

## Introduction

Studying what we cannot see, or at least what is difficult to observe, has always aroused the curiosity of human beings and fostered methodological innovations. Minute walking insects, such as micro parasitic Hymenopterans, as well as other arthropods such as acari, fall into this category. Indeed, they are difficult to observe directly in the laboratory, and almost impossible to observe in the field. Yet, they have great scientific interest for their extreme miniaturization, they are often important for the functioning of natural communities, and many have applied significance as agricultural pests or biocontrol agents. Thus, the development of new observation techniques tailored to these organisms is an important challenge^1^. This is in particular a prerequisite to access a key component of their life-history and fitness: population spread and dispersal. Dispersal is an ecological trait of paramount importance for all living organisms, as it affects not only individual fitness but also population genetics, population dynamics, and ultimately species distribution^2,3^. In biological control, dispersal is often mentioned as a key element ^4–9^. Indeed, among the characteristics which make a biological control agent a good one, there is the high searching ability, directly linked to a great dispersal capacity.

Despite its undisputed importance, dispersal remains an ambiguous term^10,11^. The simplest ones broadly refer to the increase of the distance between individuals of a group or population^12^, whilst others are more restrictive and refer to settlement and reproduction, excluding forms of movements related to foraging or seasonal migrations^13^. We here address both sorts of definitions: the spatial spread of individuals (dispersal *sensu lato*), that can have consequences on reproduction and gene flow (dispersal *sensu stricto*).

For minute insect, the relevant scale for the study of such dispersal is that of a few meters, which can be characterized both in the field and with adapted laboratory devices.

An important group of minute insects are parasitic micro-wasps, most of them egg parasitoids, i.e. insects that lay their eggs inside the eggs of other species^14^. For instance, the egg parasitoids *Trichogramma* sp., which are used in this article as a model system, are among the most commonly used biocontrol agents. These are micro-hymenopterans that are less than 0.5mm long^15^, and that mostly move by walking. Their ease of rearing on alternative hosts, and their wide geographic distribution make them useful biological control agents^16^. They are marketed by several companies and are the most widely used agents against certain pests such as the corn borer^17^. For these insects, host-seeking and parasitism are the prime activities of the adult stage, whereas pre-imaginal development occurs entirely within the parasitized host eggs, and thus entails no movement. This makes foraging movements and dispersal intimately related, and also directly connected to pest suppression performance, i.e. applied value in terms of biocontrol^18^. Indeed, from a biological control perspective, the ability to disperse not only ensures the proper distribution of insects in crops, but can also be used to reduce the workload, by diminishing the number of release points per unit area^19,20^. Studying the dispersal of these insects also gives insights into their population dynamics, which is also of great interest in the context of biological control^21,22^.

The dispersal of such minute insects is classically studied at two extremely different scales: either directly in the field, or on the contrary in very small laboratory arenas^23^. Field studies have the advantage of being very close to the real living conditions of these insects. However, the small size of these insects makes their observation in the field almost impossible, and field releases are time consuming, expensive, and yield highly variable results, all this compromises experimental throughput and reproducibility. At the other end, dispersal can also be studied by small-scale behavioural tests. Those have the advantage of offering the possibility of experimenting with speed and simplicity, allowing large numbers of replicates to be obtained. Their controlled environment, although far from natural conditions, allows good reproducibility and power.

However, they neglect important larger-scale processes such as group dynamics and long-term behavioral processes. As a result, proper evaluation of dispersal metrics is often complicated, or even lacking, in studies of these insects and, in particular, in biocontrol breeding programs^24^. That is the reason why bridging movement and population scales represents an important goal to achieve to improve the understanding and prediction of population dynamics^23^.

Two methodological developments pave the way for progress on this front. On the one hand, the development of new sensors and computational technologies led to the birth of ethomics and the broad use of ethoscopes^25^. Ethoscopes are computerized video tracking systems, simply organized with an enclosure that provides lighting and support, a camera, and an arena in which insects are released. Current ethoscopes are increasingly used for larger organisms such as *Drosophila* and mosquitoes^26^, but their arenas are of a size that do not allow access to dispersal over meaningful distances^27^. On the other hand, systems allowing the tracking of individuals directly on large scales, such as the Vertical Looking Radar, the harmonic radars or gps systems, are becoming more and more powerful^1^. But all these methods imply the installation of a sensor with properties (weight, balance, size, drag) that can affect different life history traits of the organisms (e.g. energy, movement, foraging, mating)^28^. Even if they now reach very small sizes, as the scale of organisms such as birds, monkey-fishes and large insects^29^, they are still way too big and heavy to be of any use for minute insects. The often-used ‘5% rule’^30^, according to which the device should not exceed 5% of the animal mass, cannot be respected, as these insects typically weigh less than one milligram.

To address these issues and help bridge the gap between small and large scales in studies of dispersal, we introduce here a new method extending the ethoscope concept, with two innovations: an original shape dedicated to the study of dispersal, and the use of a hi-resolution imaging pipeline. We call this system the double-spiral maze (Fig.1), and it allows the study of spatial propagation at relevant scales, while maintaining high throughput, experimental control, and low cost.

**Figure 1:**
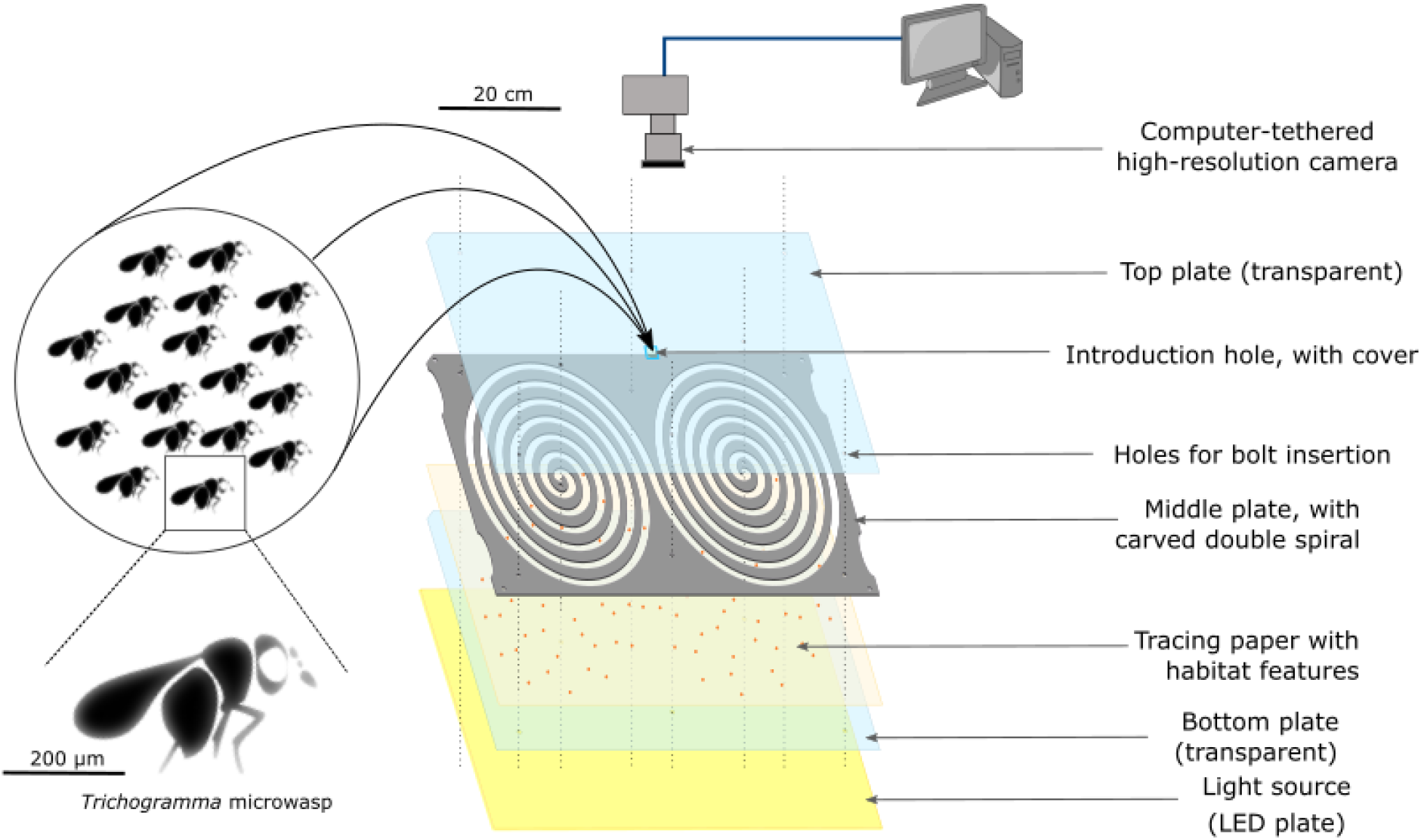
Scheme of the double spiral maze (levogyrous example). Here, the double spiral maze is included in an experimental device equipped with a LED plate, a high-resolution camera and a tracing paper with habitat features, in this case host eggs of *E. kuehniella*. The high resolution is controlled thanks to a computer.

The key concept is to wrap a long linear tunnel into a double spiral shape. This has several advantages: (I) the long tunnel can fit in a small rectangular surface, so that it is easy to locate in an experimental chamber; (II) the whole surface can be filmed efficiently with standard camera sensors, since it has similar aspect ratio, whereas a straight tunnel could not be covered with one sensor and many pixels would be lost; (III) the spiralling path ensures that any potential anisotropy in the experimental conditions (lighting, elevation, temperature…) does not correlate with distance from the centre, but rather is turned into periodic noise; (IV) the spiral path ensures that there is no sharp turns or abrupt changes that would affect movement, but instead a small gradual turning angle. This improves on previous mazes that use a zig-zag pattern^31^. Note that these advantages could be achieved with a simple spiral path, but the double spiral design brings key additional advantages: (I) the center of the tunnel, where the dispersing individuals are introduced, is located in the middle of the set-up; and (II) the path is entirely symmetrical on both sides from the introduction point: individuals dispersing to the right or to the left experience the same topology (same direction of turn, same curvatures, and same location with respect to the entire set-up).

We describe an application of the method tailored to a species of agronomic interest: *Trichogramma* parasitic microwasps^20^. The proposed system is an affordable, scalable, and easy to implement tool for insect research and biocontrol studies. In our case, the cost of making a double spiral maze and investing in a camera to capture images is just under 1434€ (see Supp.Info). In this example, a 5.75m long corridor (1cm wide and 0.5mm high) was wrapped into a rectangle of only 40cm by 30cm (see Methods “Presentation of the double spiral maze”). This makes it easy to place several copies in a standard experimental room, and to film the entire length of the corridor using one high-resolution photo sensor, allowing the detection of (very) small insects (less than 0.5mm in our case). The design of such a double spiral maze is easy with open source vector graphics softwares such as Inkscape. Once designed, it can be carved into a plate of desired thickness at low cost, for example using hot-wire cutting of Styrofoam (Burte et al. 2022) or, the solution we recommend here, laser cutting of PMMA plates.

High-resolution images of the entire maze are taken every minute for several hours, and the provided image analysis pipeline detects the location of each individual at each time point. Subsequent coordinate transformations and analyzes (see Fig. 2; and Methods) yield automated and accurate estimates of diffusion coefficients and dispersal kernels^32,33^. An original skeleton fragmentation algorithm allows to compute for each individual the linear distance from the center of the tunnel, for arbitrary shapes of the tunnel (Fig. 2 and 3).

**Figure 2:**
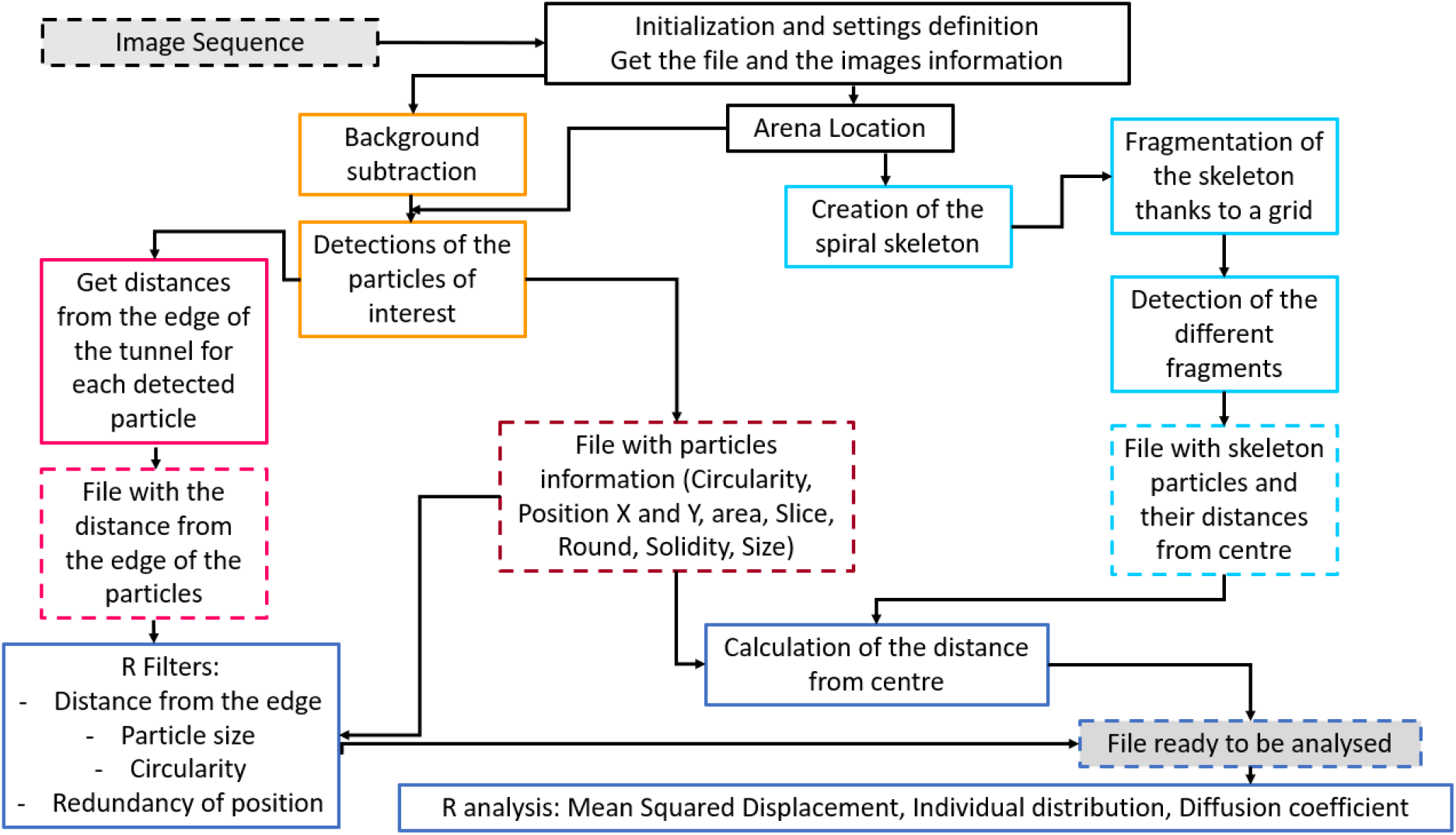
Diagram of the functioning of the automatic pipeline as well as the main steps of data processing using R software. In black, the steps related to the initialization of the file processing with all the experimental images. In light blue, all the steps corresponding to the creation and processing of the spiral skeleton. In pink, the steps concerning the distance to the edge of the individuals. In dark blue, the processing steps with the R software. The dotted line corresponds to a file. The grey boxes with dotted lines correspond to the very first file before any analysis (“Image Sequence”) and the final clean file ready to be analyzed with calculations (“File ready to be analyzed”).

**Figure 3:**
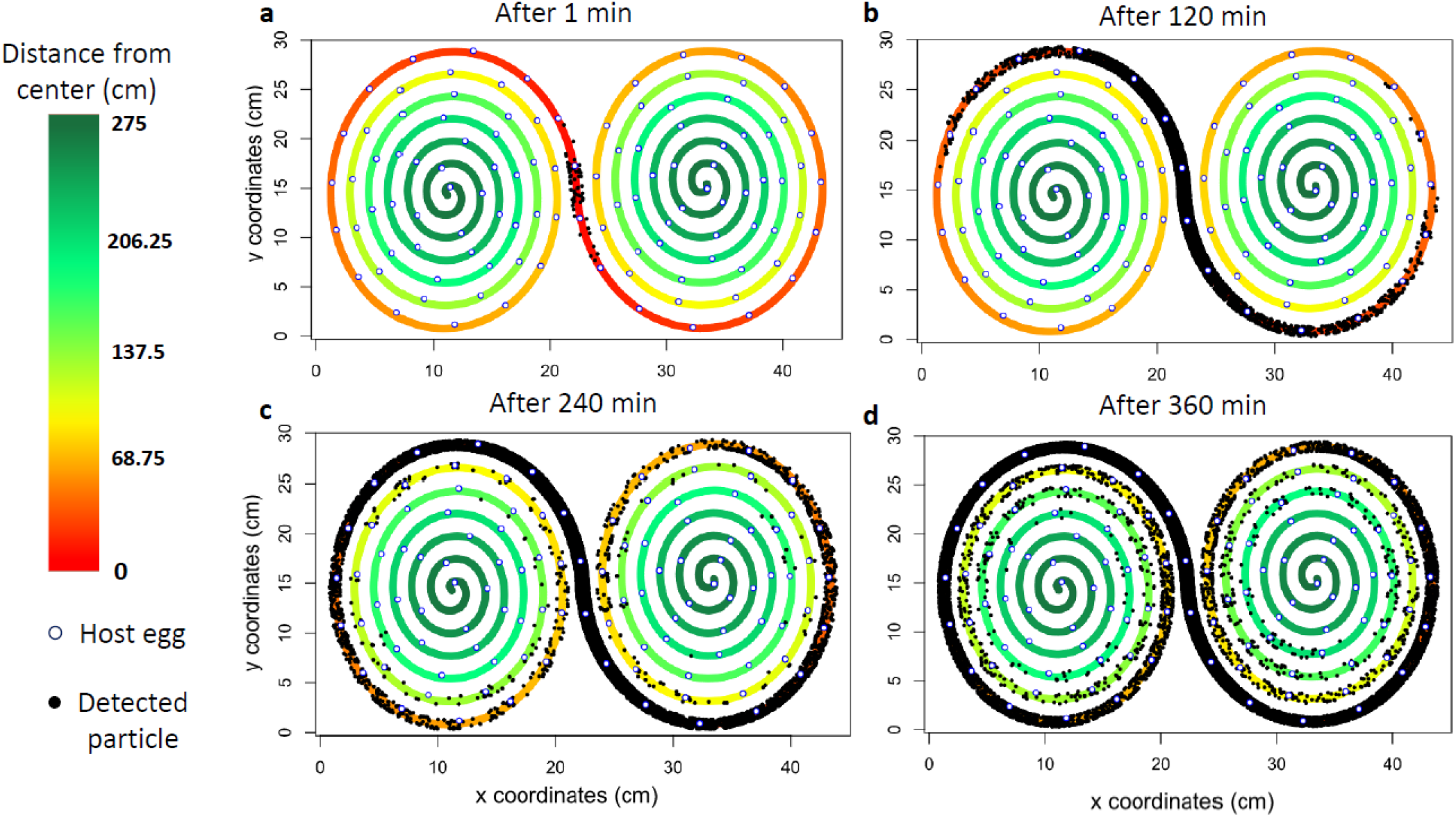
Detection of *Trichogramma* individuals at different times. a: after 1 minute; b: after 120 minutes; c: after 240 minutes and d: after 360 minutes; along the spiral path, application of our image analysis pipeline. *Trichogramma* are shown as black dots, while host eggs are shown as blue circles, filled with white.

The entire sequence of analyses can be automated thanks to an ImageJ/Fiji macro and an ImageJ/Fiji plugin, available at https://forgemia.inra.fr/ludovic.mailleret/double_maze_arena. Filtering of the outputs can be conducted with an R script available at the same address.

## Results

### Validation of the double spiral maze

#### Quality of sealing

With such a device it is essential to make sure that insects have no choice but to disperse along the tunnel and no insect can ever cut across between adjacent spires. With minute organisms, this is not an easy task, especially so when the spacing with adjacent turns is short (1 cm in our test case). To test for this, a variant of the maze was engineered with solid walls, or septa, along the double spiral tunnel, delineating 39 zones (see the Supplementary Material section “Quality of sealing” Fig.S 1). Several tests were carried out during which insects were placed at high density (about a hundred individuals) in some zones, whereas the remaining zones were left free of insects but filled with about 10 eggs of *E. kuehniella* (the substitute host). These host eggs served as sentinels for a possible leakage of insects between zones: any parasitized egg would be a sign of leakage. Each test lasted 48 hours, and direct eye checks were also carried out every 30 min for 10 hours every day. Out of 12 tests, no leakage of individuals was detected. The result tables and the detailed experimental protocol are available in the supplementary section. This confirms that the device can be sealed without resorting to glue, paraffin, or any substance between the plates, which simplifies manipulation and minimizes risks of chemical interference with insect behaviour.

#### Lighting homogeneity

Back-lighting is used to maximize contrast and facilitate insect detection^34^. A second test of importance is that of light homogeneity. Insect movement is strongly influenced by light. Tracking performance may depend on lighting parameters so that homogeneity along the entire dispersal tunnel is therefore required. To achieve this goal, the double spiral maze was backlit with a large LED plate. A pixel value profile, i.e. light intensity, was obtained on an experimental photograph to evaluate the homogeneity of the lighting. The photographs were taken with a focal length of F/22 to avoid lens vignetting, so that variations in pixel value can only be attributed to the light source. A linear scaling was established between pixel value and actual light intensity level (in lux) using standard values. We could thus establish a 2D map of the light intensity, from which we can construct horizontal and vertical light profiles, as well as the double spiral profile along the tunnel (Fig.4). The vertical profile was quite homogenous (Fig 4.b), and the horizontal profile, because of its greater length, showed some heterogeneity between the center and the extremities: the extremities were slightly less illuminated than the center (Fig 4.c). Owing to its double-spiral shape, the light profile along the tunnel was not sensitive to this long-distance trend: the average light intensity value was 9 821 lux, with no systematic variation as distance from the center increases, but rather wave-like fluctuations of more or less 14% around the average (Fig 4.d). This is a satisfying value, but of course one can further reduce these variations, if required, by purchasing a more homogeneous light source, or by applying a custom translucent mask to cancel out the heterogeneity of the source. Similarly, the value of 9 800 lux used here, which is equivalent to the light intensity in the shade in the middle of the day under a clear blue sky, can be adjusted by inflating or dimming the light source. The double spiral shape is thus able to buffer out long range lighting variations by converting them into small oscillations along the path, with no long-range trend. This is an advantage, especially since large sources of light that are perfectly homogeneous can be hard to find or quite expensive.

**Figure 4:**
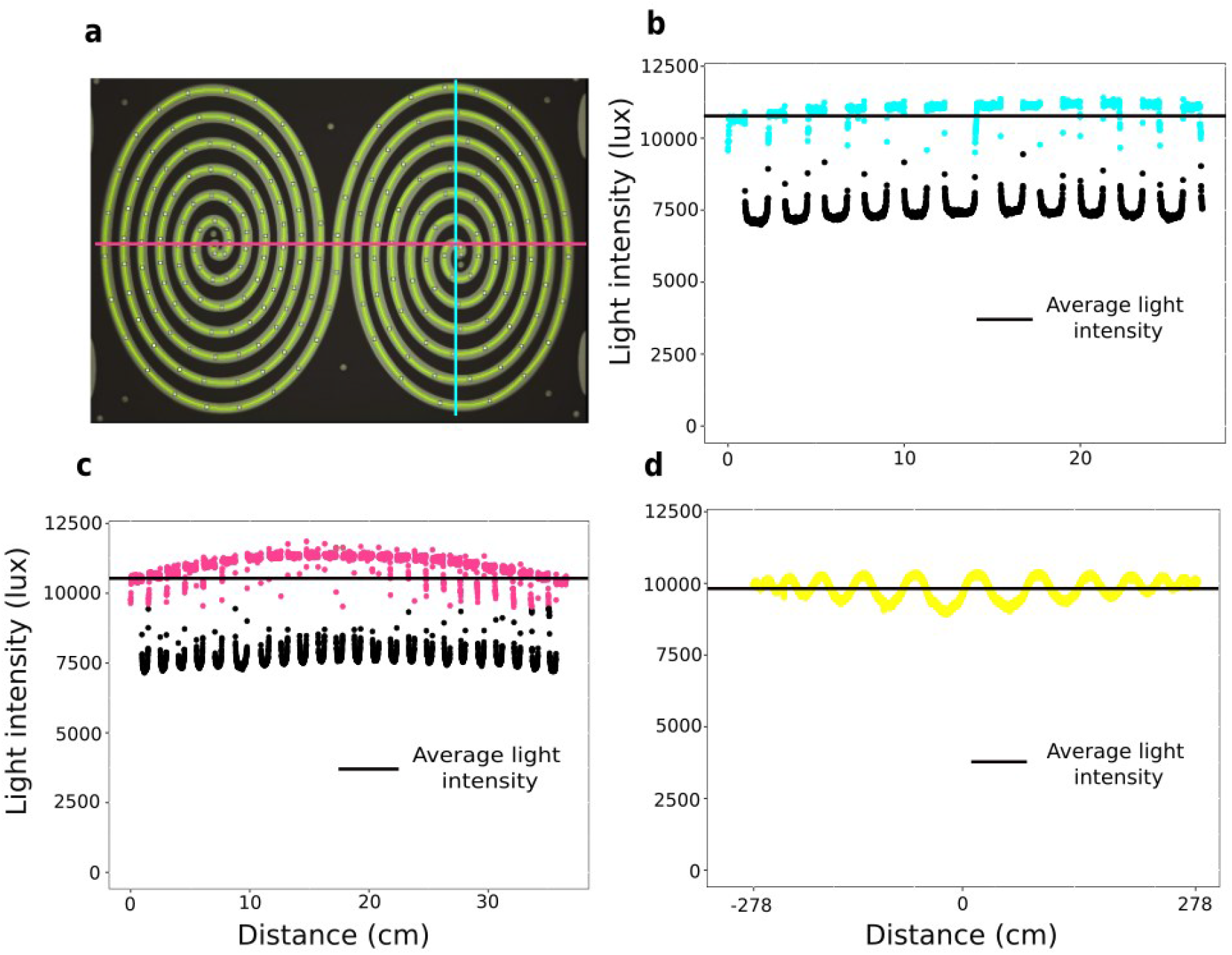
a: Schematization of the sections carried out to make the light profiles. In blue the vertical profile, in green the horizontal profile. In yellow, the path along the double spiral. b: Vertical light intensity profile. c: Horizontal light intensity profile. d: Light intensity profile along the spiral path.

#### Symmetry and spiralling direction

We checked whether the shape of the maze, i.e. the rotation direction (levo-versus dextrogyrous mazes) and the symmetry (left versus right arms) did not bias dispersal in some way. We conducted anova and skewness tests, on respectively the Mean Squared Displacement and individual repartition at each minute. The obtained p-values have been adjusted with the Bonferroni correction (see the Methods section). For both the rotation direction and the symmetry, no significant p-values were found. Therefore, rotation direction has no detectable effect on the individual dispersal inside the double spiral and no significant asymmetry could be detected in the dispersal distributions with respect to the point of introduction. To this observation, we can add that if one makes an analogy between the curvature of the walls of the double spiral and the curvature of the surface of the Earth then for the *Trichogramma* individuals the walls of the spiral are thus “curved” than the surface of the Earth is on a distance of 900 km for a human being (see the Supplementary section “Curvature of the spiral tunnel for a *Trichogramma* individual). Therefore, it is very likely that the double spiral design does not influence the dispersal of *Trichogramma* individuals.

#### Introduction of individuals

The introduction tests were carried out with the aim of achieving swift introductions of given densities of individuals (between 150 and 200). These tests showed that it was possible to easily introduce parasitoid individuals rapidly (less than 10s) and in numbers close to the target density. The detailed protocol for these tests and the table of results are available in the supplementary section.

### Validation of the hi-resolution image analysis pipeline

#### Image coverage

A comparison of the actual ground surface of the spiral with what is recorded on a photo has been performed. The double spiral has an inner surface of 504.36 cm^2^. In a photo, due to the height of the walls of the spiral (5mm), and the combination between focal length (60mm) and camera positioning (70cm) part of the floor area remains hidden and not recorded in the photo. Under the chosen experimental conditions, the surface of the floor captured in a photography was 492,64 cm^2^, therefore the lost surface was only 3%, meaning that 97% of the device surface was actually monitored with photographic captures (see the Methods section “Steps for the verification of the image coverage”). Furthermore, image coverage was not correlated with the location along the tunnel, as measured as the distance from the point of introduction of the insects.

#### Detection of individuals

Manual detection was compared to the automatic detection on ten replicates. This was done in ImageJ (see the Methods section “Manual scoring for the validation of the hi-resolution image analysis pipeline). From the ground truth data, an average of 253 individuals was observed at any given time in a particular replicate. The average number of individuals introduced in these replicates was 256, suggesting we could see about 99% of individuals. The remaining 1% were presumably overlooked (even by eye, spotting Trichogramma is not easy at that scale) or were concealed behind wall sections (and thus absent from the image data). This suggests that there is almost no loss of information between reality and the image before analysis. From the image analysis pipeline, we recovered 1817 detections, whereas the ground truth data yielded 2535 detections (Fig.5a). This corresponds to an overall 72% detection rate. The detection rate is a bit variable across replicates ranging from 68% to 78%, but stable over time (Fig. 5b). Finally, this proportion shows little variability along the double spiral tunnel except quite far from the introduction point but still non-significant (Fig.5c).

**Figure 5:**
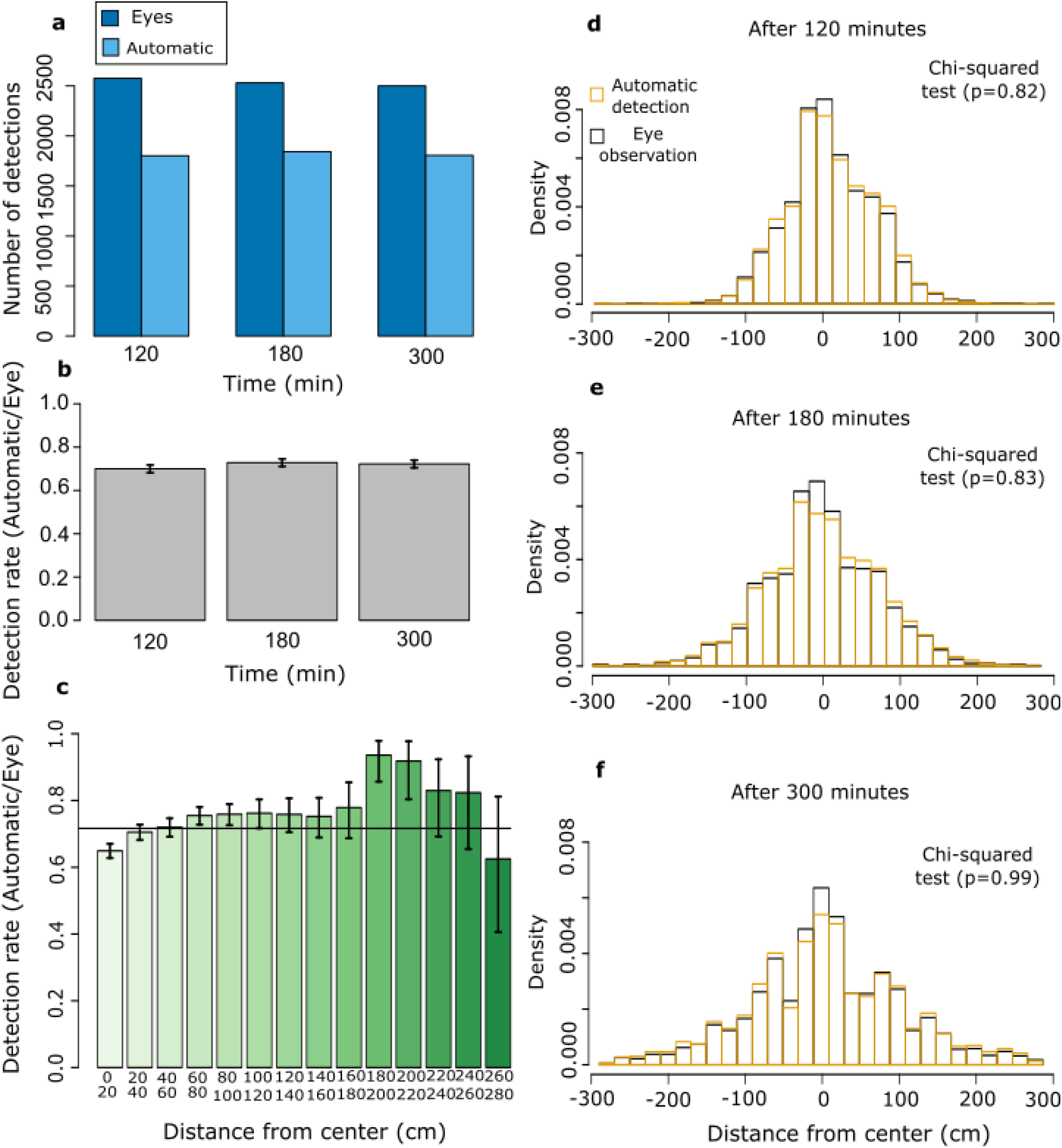
Left column: detection rates. a: Evolution of eye and automatic detection over time. b: Evolution of the detection rate over time. c: Evolution of the detection rate along the path of the spiral. Times are pooled. Here, the right and left sides of the spiral have not been distinguished but pooled, hence the graduation from 0 to 280. Right column: population distributions over space and time. In black, the distributions obtained with the eye’s observations. In orange, the distributions obtained with the automatic pipeline. d: Spatial distributions after two hours. The two distributions are not significantly different (Chi-squared test ; p=0.82). e: Spatial distributions after three hours. The two distributions do not significantly differ (Chi-squared test ; p=0.83). f: Spatial distributions after five hours. The two distributions do not differ (Chi-squared test ; p=0.99).

#### Spatial distribution of individuals

The automatic pipeline must be reliable in measuring the distribution of individuals, an important metric in dispersal analysis. For the ten replicates, the true distribution and the one inferred from image analysis were very similar. If we look in greater detail, at particular times, the same conclusions hold. Indeed, the two distributions were not significantly different at any time (Chi-square tests; Fig. 5.D.E.F).

### Data-storage requirements and data-processing times

For a 6-hour experiment with one replicate, a minimum of 70 GB of storage space is required. The 360 photos in RAW format represent 30 GB. These photos must then be converted into TIFF format to be analyzed by the pipeline, which represents 40 GB for a transient file. The conversion of the photographs and the application of the pipeline each requires 45 minutes.

### Example of practical application: is dispersal by groups of *Trichogramma* diffusive?

The double spiral maze and the accompanying automatic detection method can answer several questions of ecological and agricultural interest. We were interested in assessing intraspecific variation in dispersal dynamics within species of *Trichogramma cacoeciae*. A way of understanding spatial spread is to look at its diffusivity. If we assume that *Trichogramma* individuals’ movement is random, i.e. if individuals move independently in all directions without preference, at the relevant scale, a standard diffusive dispersal is expected^16^. To check whether the observed dispersal is diffusive or not, it is possible to look at the Mean Squared Displacement (MSD). For a group of individuals initially introduced at the same location, the MSD corresponds to the variance of the distribution of individuals at a given time. It should increase linearly with time in the case of a simple diffusion^32^.

We conducted a screening of 14 strains of *T. cacoeciae* (see the Supplementary Material section “Strains information”), all sampled in France using *E. kuehniella* sentinel eggs and maintained in the laboratory under identical conditions. We evaluated each strain for dispersal with 10 replicates. In each replicate, we released 200 females on average, all of the same age, in the double spiral maze, between 8 and 8:30 am. One *E. kuehniella* host egg was glued every five centimetres along the double-spiral tunnel. We stopped the experiments after six hours and took high resolution images every minute and analyzed them with the above-described pipeline. Thanks to the size of the maze, four strains could be tested in parallel every day, in one room.

By applying the image analysis pipeline, we could easily compute the Mean Squared Displacement that each strain had achieved in six hours, and thus rank species in terms of their dispersal propensity (Fig. 6a). We regressed MSD over time with simple or piecewise regressions, for each of the 14 strains (see Methods section “piecewise linear regression”). Overall, there was significant inter-strain variation in MSD, revealing quantitative genetic variation for this phenotype: 33% of total MSD variation was attributed to the strain effect (see Methods section “piecewise linear regression”) (Fig. 6a). More interestingly, we could discover inter-strain qualitative differences in the dynamics of dispersal. Moreover, from these analyzes, we identified three distinct types of dispersal dynamics (Fig. 6). The first one (type 1) corresponds to strictly diffusive spatial spread, as illustrated by strain ISA1075 in Fig. 6b. However, only 14% (2/14) of the tested strains possessed this behaviour; all other strains showed a non-diffusive dispersal, which can be classified into two categories: type 2 and type 3. Type 2 is characterized by a latency phase with relatively slow spatial spread, followed by a fast phase with sustained faster spatial spread. This is illustrated for strain ACJY144 in Fig. 6c, for which the latency phase lasted about two hours. This result is consistent with the fact that the distribution of ACJY144 individuals is not normal since for simple diffusive movement the distribution of individuals at a given moment in time is a Gaussian (Fig.6). Type 3 was the most frequent type of dispersal (7/14 strains). It is characterized by a latency phase, followed by a fast phase, but eventually the rate of spatial spread slows down again. This is illustrated for strain TCMz in Fig. 6d, for which the slow phase occurred after about four hours. A couple of strains (“Others’’ in Fig. 6e) had non-diffusive dispersal as well, with three distinct phases (as in type 3), but specific patterns (see the Supplementary Material section “Example of practical application: description of the category “Others” “ Fig. S2). Differences in types of dispersal were independent from differences in total MSD (p=0,19; Supp. Material Fig. S3). This revealed that the vast majority of strains has more complex dispersal than the standard diffusion model commonly assumed, even in simple experimental environments.

**Figure 6:**
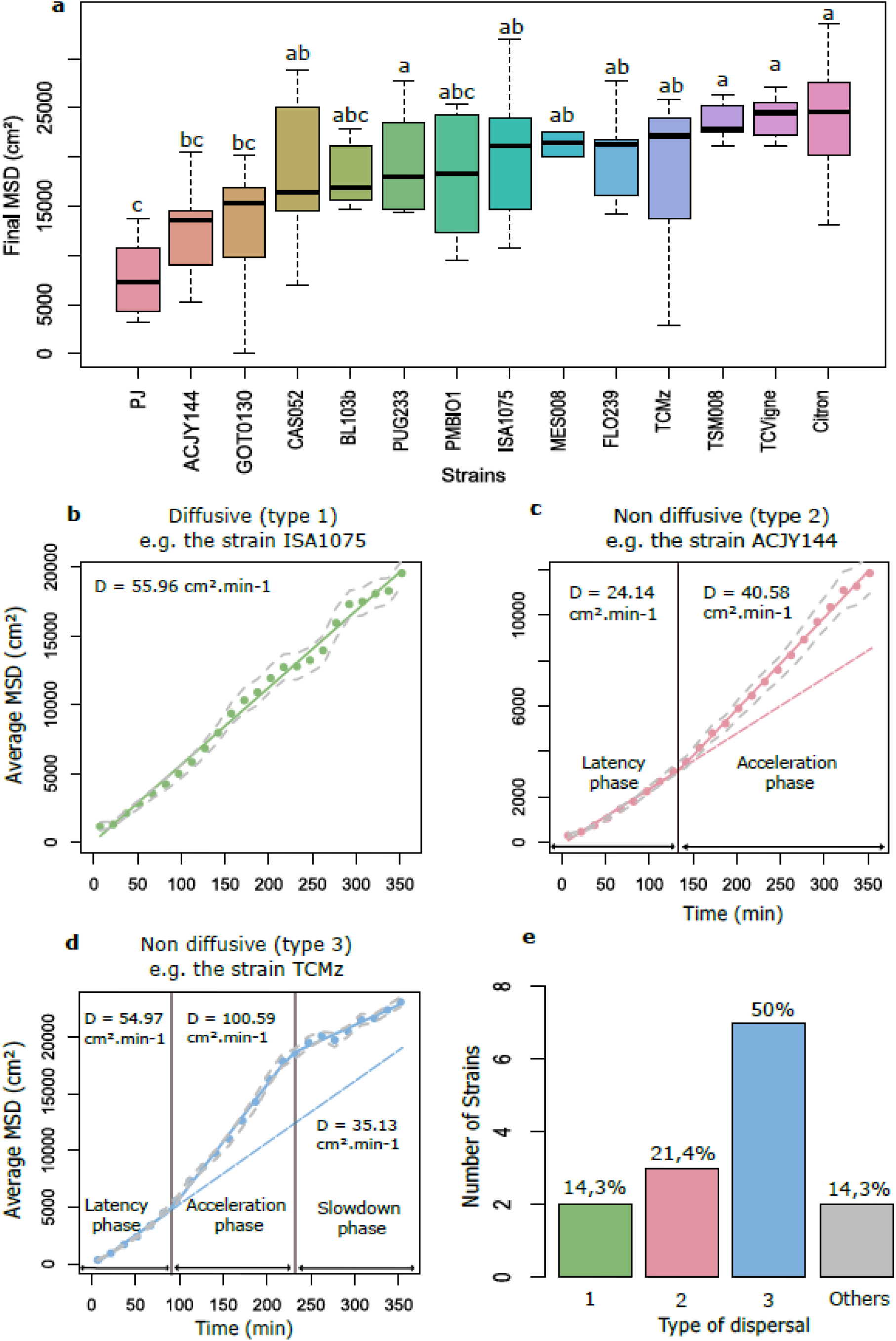
(a) Boxplot of the final Mean Squared Displacement for all tested strains. Letters indicate significant differences as obtained from Tukey’s post hoc anova tests. Plots representing the Mean Squared Displacement over time for strain ISA1075 (b), representing diffusive (linear) dispersal (type 1); strain ACJY144 (c) representing non diffusive dispersal with two phases (type 2); strain TCMz (d) representing non diffusive dispersal with three phases (type 3). The figure (e) represents the percentage of strains belonging to each dispersal type. In panels B-D, the coloured dots represent the raw data averaged for each time window. The solid lines correspond to the best supported piecewise regression models. The grey dashed lines correspond to the confidence intervals calculated on the MSD means obtained by replicates. These intervals therefore account for inter-replicate variability. The black vertical lines indicate the breakpoints identified by the piecewise linear models. The dashed lines of the same colour as the regression sections correspond to the extrapolation of the linear regression of the first phase.

## Discussion

We developed a double spiral maze with the goal of quantifying minute insects’ dispersal metrics at spatial and temporal scales similar to those that are relevant in the field (several meters, several hours), while retaining the experimental power and throughput of ethoscopes. Indeed, currently there exists a lot of methods to monitor the movements of insects, e.g. through the use of radars, traps, or even eye observations, but those are only applicable with organisms big enough^1^. For smaller organisms such as *Trichogramma* parasitic wasps and similar minute insects, all these techniques fall short. As a result, their movement is often only assessed at two extreme scales: either indirectly with field trials or directly with small laboratory arenas. The former is time consuming and returns small amounts of data with considerable noise especially since dispersal is very sensitive to environmental variations^1,24,35–37^, whereas the latter neglects larger scale phenomena such as group dynamics or variations in activity patterns due to different extrinsic factors^38^. This last point can be seen as a limitation but also as an advantage since detaching from variability factors allows for more repeatability. But it can make it difficult to establish a link between laboratory observations and what is actually happening in the field. Furthermore, the two approaches are so disparate that they are often hard to reconcile. This leads to a lack of robust experimental data for analysing the dispersal of minute insects. In this context, the double-spiral maze can offer a complementary, intermediate scale of study, borrowing the experimental power of microscale laboratory studies, while reaching spatial and temporal scales close to field processes. In this sense, it is a “mesoscale” set-up.

We have shown here that our system allowed to quantify dispersal metrics with great precision and identify qualitatively different types of dispersal dynamics. The method returns high quality data, with an average detection rate greater than 70%, that is stable along the path of the double spiral maze and over time. It is straightforward, with no other change in the protocol, to increase this performance if required, by increasing the frequency of image captures (e.g. one every 10s is totally feasible) and/or increasing image resolution (50-60 Mpx sensors already available). This would come at the expense of greater data-storage requirements and data-processing times but would provide even greater experimental power. The original shape of the double spiral maze brings decisive advantages, and we showed that it had no appreciable impact on the results obtained with our study system *Trichogramma cacoeciae*. Of course, such a verification should be made for any other study system, but our results suggest it would work equally with many similar organisms. We could obtain a satisfying lighting with few variations at low cost, though one could certainly reduce these variations further, if required, by purchasing more homogeneous light sources, or by applying a custom translucent mask to cancel out the heterogeneity of the source. Similarly, the light intensity of 9,800 lux used here can easily be adjusted by inflating or dimming the light source. The nature of the lighting can also be adjusted; for instance, switching to IR lighting to monitor the activity of organisms at night for example, would require only minor adjustments. The use of laser cutting allows for ease and speed of production, as well as perfect reproducibility between prototypes. The double spiral maze is therefore an inexpensive and very scalable device. The technical robustness of the double spiral maze and its versatility, a direct result of its conception, allows to raise and address a broad range of biological questions.

As an example, the double spiral maze can be used to screen strains or genotypes and to quantify intraspecific and interspecific variation in dispersal. Furthermore, the possibility to add environment features such as host eggs, as we have done, offers the possibility to study parasitism rates, relate them to individual movements, and study the effects of host density and distribution on population spread. The location of individuals also permits to identify patterns of host processing and interactions. The possibilities are rich beyond the simple application presented here. In this article, we could document intraspecific genetic variation in the final MSD after six hours, with strains dispersing significantly more (such as Citron, TCVigne, TSM008) and others being more sedentary (like PJ or ACJY144; Fig.7A). This is the first report of such variation in this species. Furthermore, we could demonstrate that most strains did not undergo simple diffusive dispersal (only 4% had) but had more complex types of spatial propagation involving two or three phases (Fig. 7). This non-linearity of the MSD is consistent with the fact that the distribution of individuals was often not normal (see e.g., ACJY144 in Fig. 6), as one would expect in the case of classical diffusion^32^. Interestingly, the type of dynamics was unrelated to the final MSD, and so strains with similar final MSD could exhibit different temporal dynamics of spread: this was the case for strains Citron (type 2) and TCVigne (type 3). The type of dynamics is thus an additional, independent, component of the dispersal phenotype, and one that could only be characterized with the continuous tracking that our imaging pipeline permits.

Several mechanisms could explain the existence of these different types of dispersal. Type 2 is characterized by a latency phase with relatively slow spatial spread, followed by a sustained phase of faster spatial spread. The latency phase can be as long as two hours and fifteen minutes (strain ACJY144). This is much longer than typical small scale and behavioural experiments, stressing the difficulty to scale-up from short-term observations and the importance of studying spatial spread at larger scales. Type 3 is also characterized by a latency phase followed by a fast phase, but eventually the rate of spatial spread slows down again after several hours. Individuals did not reach the tips of the maze in six hours: this occurred very rarely (a few individuals, in only two out of 140 replicates). Therefore, type 3 dispersal cannot be accounted for by a boundary effect. The slowdown phase may rather reflect the fact that females have already parasitized several hosts by that time, and therefore switch to a different exploration pattern^39^. A relationship between egg load and activity rate has already been described in other parasitoid species, and attributed to “ovarian pressure”^40,41^. Of course, these differences in dispersal dynamics might also incorporate potential variations in diurnal patterns of activity among strains. Elucidating the behavioural mechanisms responsible for these different types of dispersal and their genetic basis is beyond the scope of this article but opens many interesting research questions.

The double spiral maze was conceived for the particular case of minute insects since it is very difficult to monitor them in the field. However, the system can easily be adapted to other small walking organisms, such as acari, ants, aphids or carabids, or even larger organisms for which the use of the double spiral would facilitate experiments and the obtention of larger amounts of data. Indeed, its ease of design and fabrication allows for easy modification of its dimensions. There exist laser cutters that can cut dimensions as large as 3×2 meters for example^42^, so that mazes that are broader or longer (beyond 20 meters) are straightforward to build. Even more spectacular scales could be attained using other construction methods, to implement the double spiral maze with larger organisms for which dispersal is often studied, such as the common lizard^43,44^. It is also straightforward to alter the shape of the tunnel and, for instance, to engineer large sections along it, separated by narrower sections (in the vein of the approach in Fig. S1), to mimic a linear sequence of connected habitat patches. Beyond terrestrial organisms, the double spiral maze can be adapted to aquatic organisms with minor adjustments, as questions of movement and dispersal are also widely addressed in such organisms, from aquatic micro-organisms to fish^45,46^.

To conclude, we introduced a new type of ethoscope and analysis pipeline that helps bridge the gap between small and large scales in experimental studies of movement and dispersal. This opens new perspectives to extend our understanding of dispersal in different groups of living organisms. This is true in particular for very small insects, an under-studied yet important group for which other methods based on the use of sensors are not adapted^28,30^. From a more applied point of view, with organisms such as *Trichogramma* that are used in biocontrol, the double spiral maze offers the possibility of reducing greenhouse or field experiments necessary to select the species and strains, and to conduct high-throughput screenings on larger sets of candidate species and strains. Selection on searching capacity would thus not be secondary anymore^47^.

## Methods

### Micro wasps

*Trichogramma cacoeciae* individuals used in this study were supplied by the INRAE biological resource centre “EggParasitoids Collection” (CRB EP-Coll, Sophia Antipolis) which continuously rears a panel of Trichogramma strains. The strains used in these experiments were collected in France in different geographical locations, and on various plant species, between 1987 and 2016. These strains have been raised on irradiated *Ephestia khueniella* eggs, on a 14 days development cycle at 25 and 18 °C.

Approximately 200 parasitized eggs of *Ephestia khueniella* stuck on strips were isolated 36h before the experiments in small PCR tubes for each strain to be tested 36h later. A drop of honey was placed in each of these tubes. These tubes were then placed at 25 degrees so that the females would emerge and be one day old. On the day of the experiment, the strips with the eggs from which the females emerged were removed in order to allow the introduction of the individuals in the double spiral mazes. Thus, an average of 185 individuals were introduced in each replicate.

### Construction and assembly of the double spiral maze

Here we present in detail how the device is designed. It consists of three superimposed plates:

- An intermediate plate in white opaque Polymethylmethacrylate (PMMA, Plexiglas) of 440×330mm in which the double spiral has been laser cut with a laser cutting machine Trotec Speedy 100. The double spiral thus cut forms a 5,75m tunnel, 1cm wide and 0.5cm high. The height/width ratio of the device was chosen to match that of the photo sensor used (24×36mm) during the experiments, and thus maximise the use of the sensor pixels.
- Two transparent Plexiglas plates, framing the double spiral plate at the bottom and top. The top plate also has a central hole through which the individuals under study can be inserted. The three Plexiglas plates are pierced at eight points, which constitute the clamping points that hold the three plates together and ensure that the device is airtight, but above all that it is airtight to the individuals that disperse in it. The entire device is illuminated from below by a LED plate (119.5*59.5 cm), which induces a backlighting effect, thus creating a stronger contrast between the insects studied and the rest of the lighting (light tunnel of the spiral). Dextrogyre and levogyre mazes have been created in case the direction of rotation influences the observed results. No significant effect has been detected (see Methods). In the experiments we conducted, a tracing paper sheet with *E. khueniella* eggs has been added between the last PMMA plaque and the double spiral path.

It is important to note that to assemble the whole device, only three plates of PMMA, a quick and easy design with the free software Inkscape and access to a laser cutting machine are necessary.

In our case, experiments were performed using the previously described double spiral maze device and its lighting system. Individuals were introduced as described in the introduction test protocol. The mazes were placed under NIKON D810 cameras equipped with an AF-S Micro NIKKOR 60mm f/2.8G ED lens, at a height of 70cm. The aspect ratio of the device was chosen to match that of the photo sensor used (24×36mm), and thus maximize the use of the sensor pixels. The system as presented here offers a resolution of 36 Mpx.

The double spiral maze was surrounded by a box preventing stray light. This box is an empty rectangle with 4 walls covered with black canson paper, with a height of 8 cm.

The focus, camera settings adjustments, and camera trigger were made using the free *DigiCam Control 2* software, V.2.0.0. The following settings were used to optimize the exposure of the image: ISO 125, Shutter speed 1/125 and aperture f/11.0.

Once all the adjustments were made. The image capture was launched for six hours, with a photograph taken every minute. The images were saved in RAW format (for storage efficiency) and transformed to TIFF format using *Darktable* software, prior to analysis. Converted images were then analyzed with the ImageJ/R pipeline.

### Image analysis pipeline for the automatic detection of individuals through time

The image analysis pipeline can be found at https://forgemia.inra.fr/ludovic.mailleret/double_maze_arena under the name “Principal_Macro”. It can be easily run in ImageJ using the following sequence of actions:

#### Plugins>Macros>Run

Then select the right macro file to be used.

The image analysis pipeline starts by a conversion of the images into grayscale (8-bit). Then the moving particles are detected by the calculator function. A threshold is then chosen so that the moving particles detected correspond only to insects, not artifacts. The analysis of the particles is then launched, yielding the coordinates of each particle detected.

The automatic pipeline for image analysis is applied to all the images obtained during the experimentation, i.e. 360. To reduce the needs in memory, images are loaded by packs of 60 images, in a sliding window manner (adjustable parameter). Here is the main sequence of actions carried out in ImageJ by our pipeline:

- *File>Import>Image Sequence*: Importation of the image sequence to be analyzed.
- *Image>Type>8-bit*: conversion of the images in 8-bit, i.e., in shades of grey. The software assigns to each pixel a value from 0 (black) to 255 (white).
- *Image>Stacks>Z project*: tracking and analysis of the particles of an image over time to define a still background image, on which the particles move. The background image is subtracted from the actual image to distinguish the moving particles.
- *Process>Image calculator*: particles tracking on each image, based on the background calculated previously. During this process, the background is removed to leave only mobile particles (the “Subtract” option must be chosen and “32-bit (float) must be used.
- *Image>Adjust>Threshold*: Definition of the threshold on the images from which the background has been subtracted, to consider only the trichograms. Simply uncheck “Dark Background” and set the sliders so that only the trichograms are detected in red.
- *Analyze>Set Measurements*: Step to select the set of parameters that will be measured with the *Analyze particles* function. We have selected shape parameters: circularity for which a value of 1 indicates a perfect circle and 0 an elongated shape, solidity to make an area selection convex, area of the detected particle in square pixels and the aspect ratio which corresponds to the description of the ratio of the width and height of a given pixel.
- *Analyze>Analyze Particles*: Launching the analysis, based on the parameters previously set.

Upstream, the skeleton of the spiral (line tracing the center of the path of the double-spiral maze) is realized on each image sequence. The macro to realize the skeleton is available at https://forgemia.inra.fr/ludovic.mailleret/double_maze_arena under the name “Macro_Skeleton” and needs this installation of the plugin “Spi_rou” available at the same address. The skeleton is realized on the calculated background:

#### Process>Binary>Skeletonize

This “skeletonize” is used to create a skeleton along the double spiral path. The resulting skeleton is then cut into small particles using a grid that creates a cut point at each meeting point between it and the skeleton to create a particle. The center of the spiral is then defined by drawing the diagonals of the spiral. It is therefore possible to obtain for each particle its position relative to the center, giving us the possibility to access its distance to the center. Later, in R, we will be able to give a position to each detected particle by associating its distance to the associated skeleton particle thanks to an orthogonal projection.

The last step of this image analysis pipeline is realised with the R software. The data obtained from the automatic particle detection are cleaned thanks to different filters to obtain particles which only correspond to *Trichogramma* individuals (Figure 4):

- A filter on redundancy, assuming that one trichogramma cannot be twice at the same spot over time.
- A filter on particle size, knowing the average size of a trichogramma individual.
- A filter on their distance from the edge of the spiral path.
- A filter on the shape of the particles detected using a circularity value, knowing that trichogramma individuals are quite circular particles.

### Validation of distribution symmetry and effect of rotation direction

To check whether the shape of the maze, i.e., the rotation direction (levo-versus dextrogyrous mazes) and the symmetry (left versus right arms) did not bias dispersal in some way. We conducted tests on 14 strains of *Trichogramma cacoeciae*, at each minute of time. Therefore, to all the p-values obtained for both rotation direction and symmetry, a Bonferroni correction has been applied. For the rotation direction, anova models have been run on simple linear models, with the Mean Squared Displacement as the explained variable and the rotation direction as the explanatory variable. For the symmetry of the repartition of the individuals inside the maze, the “skewness.norm.test” function which compares the skewness of the distribution to the theoretical value of 0, has been used.

### Steps for the verification of image coverage

In order to compare the real surface of the double spiral maze path with the surface of what is actually photographed, we performed a calculation of the surface of the double spiral path on a photograph using Image J software, as well as an area calculation on the spiral cutting drawing in Inkscape software.

In the ImageJ software, after having calibrated the scale in cm, we thresholded the image using the “Color Threshold” tool in order to obtain a thresholding selecting only the interior of the spiral path. Then we created a binary version of the image. Finally, using the “Analyze particles” function, we could obtain the area of the thresholded part.

At the same time, with the Inkscape software, we simply selected the interior of the spiral path and with the menu Extension, Visualize path, Measure path, we selected as the variable to be measured the area.

The two areas thus obtained were compared using a ratio.

### Manual scoring for the validation of the hi-resolution image analysis pipeline

To assess the performance of the automatic detection of individuals, we performed a manual scoring of individuals present in the maze on ten replicates at three different times: after two, three and five hours. After this, the location of all tags on all images was automatically recovered using an ImageJ macro. The manually annotated (x,y) coordinates of *Trichogramma* individuals were transformed into « distance from the introduction hole » by projecting them onto the skeleton of the spiral as previously explained. This provided the ground truth for this set of images, which could be compared to the corresponding results obtained using the automatic image analysis pipeline.

### Piecewise linear regressions and statistical tests

To ensure data independence, MSDs measured at each minute were averaged across replicates over 15 minutes time windows. Therefore, instead of having 360 measurement points, 24 averaged measurements remained. The time window of 15 minutes was chosen after previously testing different times: 2, 5 and 10 minutes. The autocorrelation between successive residuals (lag 1) was checked with the *acf* function applied on the residuals of each section. The time window was extended until all lag1 autocorrelation was no longer significant. With 10 minutes, there was still 23% autocorrelation.

Once the time window was chosen, averages on the time window were calculated for each strain and replicate and finally averaged to have one value per 15 minutes for each strain. On these data, simple regressions, with the *lm* function, as well as piecewise regressions with two or three pieces, with the segmented function from the package segmented, were generated for each strain. Likelihood Ratio Tests were performed with the *anova* function, to assign the best regression to each strain.

By obtaining these data we were then able to investigate how much of the variability in final MSD was explained by differences between strains. To do so, an anova model was performed on the final MSD with the following explanatory variable: Replicate which accounts for the variability associated with the days of experience; the number of detections by the pipeline for each replicate, the orientation of the double spiral maze (levogyrous or dextrogyrous) and finally the strain. As outputs, we recovered the sum of squares of the treatment “Strain” (SST(1)) and the sum of squares of the residual (SSR) error, as well as the sum of squares of the other treatments (SST(2)) to calculate the following ratio: SST(1)/(SSR+SST(1)+SST(2)).

## Code availability

Code to use the image analysis pipeline and to filter the obtained data is found at https://forgemia.inra.fr/ludovic.mailleret/double_maze_arena with a Gnu general public license.

## Acknowledgments

We warmly thank all the members of the “SoFAB” (the FabLab of Sophia Antipolis) who helped us to develop the double spiral maze by sharing with us their knowledge and their know-how with the laser cutting machine. Fundings are from INRAE (SPE Department) and Université Côte d’Azur (IDEX UCA-JEDI) which supported Mélina COINTE’s PhD.

## Authors’ contributions

MC, VC, LM, GP and VB designed the experiments. MC and GP performed the experiments. MC and VC analysed the data. MC and VC wrote the manuscript with inputs from all authors.

## Supplementary Material

**Table 1:** Evaluation of the cost of production and use of the double spiral maze. The dimensions of the sold plates actually allow the production of two double spiral mazes.

| Material | Number of units | Unit price (€) | Total price (€) | Commercial reference |
| --- | --- | --- | --- | --- |
| Opaque Plexiglas plate of 1000x1000 mm | 1 | 34.4 | 34.4 | Richardson |
| Transparent Plexiglas plate of 1000x1000 mm | 2 | 40.95 | 40.95 | Richardson |
| Camera Nikon D810 | 1 | 1298 | 1298 | Idealo.fr |
| Use of a LASER cutting machine hourly rate | 1 | 60 | 60 | SoFab Sophia Antipolis |
|  |  |  | **1433.4** |  |

**Table 2:** Results of the quality of sealing test

| Strain | Test Duration (hours) | Mean number of individuals in the tested zones | Leaks observed by eye | Parasitized sentinel eggs | Tested zones |
| --- | --- | --- | --- | --- | --- |
| PMBIO1 | 48 | 78 | None | 0 | 17,16,11,19,31,27,37 |
| PMBIO1 | 48 | 46 | None | 0 | 6,8,16,17,19,22,26,37 |
| ACJY144 | 48 | 128 | None | 0 | 1,3,13,21,25,29,39 |
| PMBIO1 | 48 | 71 | None | 0 | 10,2,35,28 |
| PJ | 48 | 60.5 | None | 0 | 5,15,7,30,36,20 |
| PUG029 | 48 | 64.2 | None | 0 | 32,38,24,12,9,14 |
| PUG029 | 48 | 73 | None | 0 | 17,16,11,19,27,34 |
| PMBIO1 | 48 | 61 | None | 0 | 6,8,22,37,31 |
| ACJY144 | 48 | 44.4 | None | 0 | 33,1,3,13,21,25,23,39 |
| PMBIO1 | 48 | 65,2 | None | 0 | 10,2,35,28,15 |
| PMBIO1 | 48 | 86 | None | 0 | 5,15,7,30,36,20 |
| PMBIO1 | 48 | 76.5 | None | 0 | 32,38,24,12,9,14 |

**Table 3:** Results of the introduction of individuals test

| Strain | Introduced<br>Individuals | Remaining<br>individuals<br>inside the tube | Total of<br>individuals | Number of eggs<br>on a strip<br>introduced in<br>the tube | Introduction<br>Rate |
| --- | --- | --- | --- | --- | --- |
| PMBIO1 | 45 | 38 | 83 | 100 | 54.2 |
| PMBIO1 | 78 | 51 | 129 | 150 | 60.5 |
| ACJY144 | 99 | 60 | 159 | 220 | 62.3 |
| PJ | 109 | 45 | 154 | 600 | 70.8 |
| PMBIO1 | 166 | 48 | 214 | 500 | 77.6 |
| PMBIO1 | 96 | 60 | 156 | 200 | 61.5 |
| PMBIO1 | 92 | 65 | 157 | 200 | 58.6 |
| PMBIO1 | 63 | 27 | 90 | 200 | 70 |
| PMBIO1 | 62 | 24 | 86 | 200 | 72.1 |
| PMBIO1 | 92 | 70 | 162 | 200 | 56.8 |
| PMBIO1 | 90 | 73 | 163 | 200 | 55.2 |
| PMBIO1 | 80 | 40 | 120 | 200 | 66.7 |
| PMBIO1 | 79 | 30 | 109 | 200 | 72.5 |

**Table 4:** Information on the 14 strains used in the results section: their name, their sampling location, the plant on which they were captured and their sampling year.

| Strain | Sample Location | Capture Plant | Sampling year |
| --- | --- | --- | --- |
| **ACJY144** | Aime, France | Ash tree | 2015 |
| **BL103b** | Le Change | Primrose | 2016 |
| **CAS052** | Castellane | Boxwood | 2016 |
| **Citron** | Grasse | Lemon tree | 2012 |
| **FLO239** | Le Crouzet | Rose bush | 2016 |
| **GOT0130** | Gotheron | Apple tree | 2016 |
| **ISA1075** | Maurice de Beynost | Plum tree | 2015 |
| **MES008** | Beaumat | Maple tree | 1987 |
| **PJ** | France | Laboratory strain | 2015 |
| **PMBIO1** | France | Unknown | 2014 |
| **PUG233** | La Pugère | Apple tree | 2015 |
| **TCMz** | France | Unknown | 1987 |
| **TCVigne** | France | Unknown | 1987 |
| **TSM008** | Tours sur Meymont | Apple tree | 2015 |

**Figure S1:**
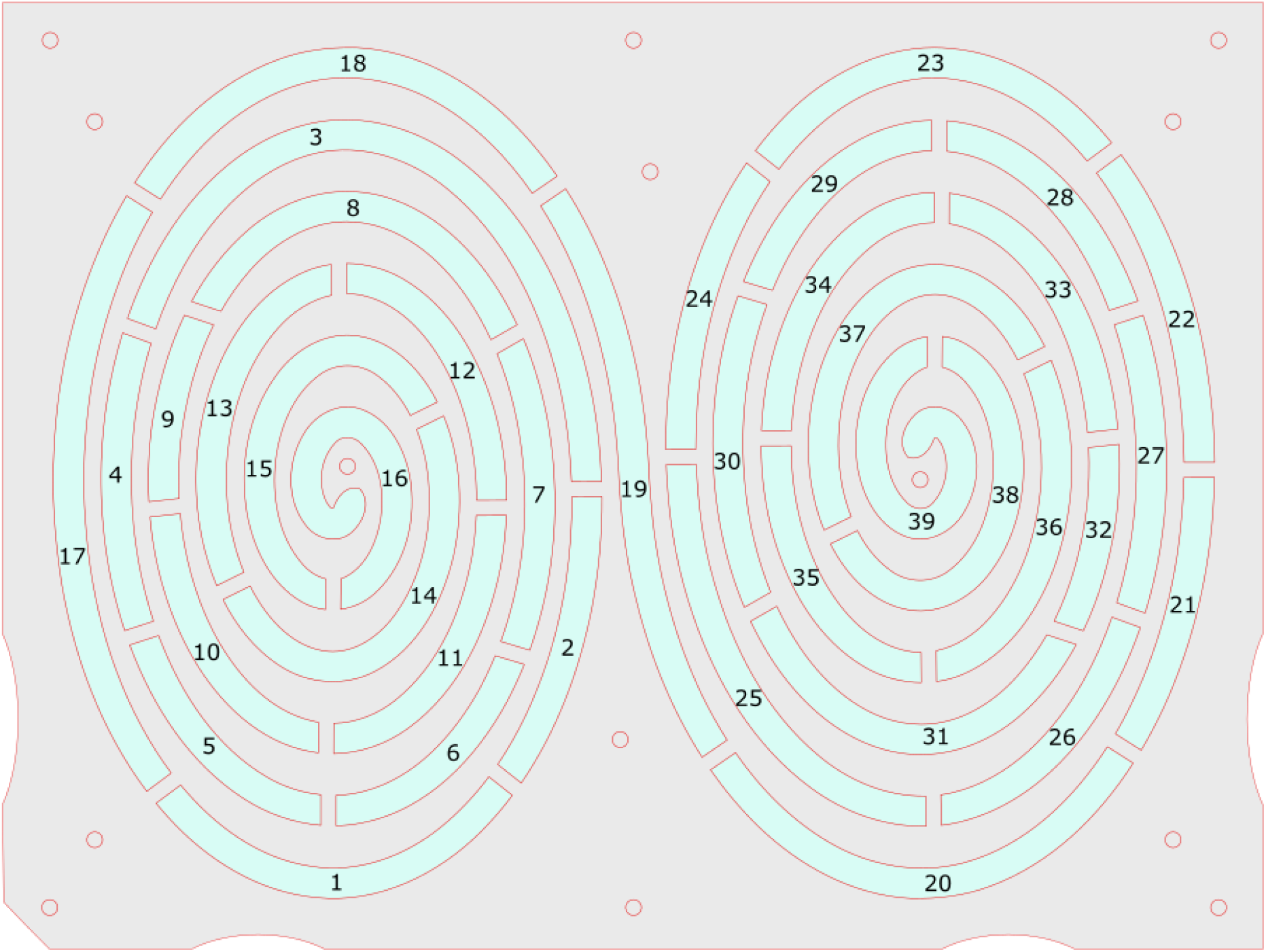
A variant of the (levogyrous) double-spiral maze that was engineered with solid walls (5mm width), or septa, along the tunnel, in order to test for the quality of sealing. This was designed with the Inkscape software, delineating 39 isolated zones (in light blue on the scheme). The numbers correspond to the zones called in table 2. An SVG file ready for laser-cutting is available as Supplementary File.

**Figure S2:**
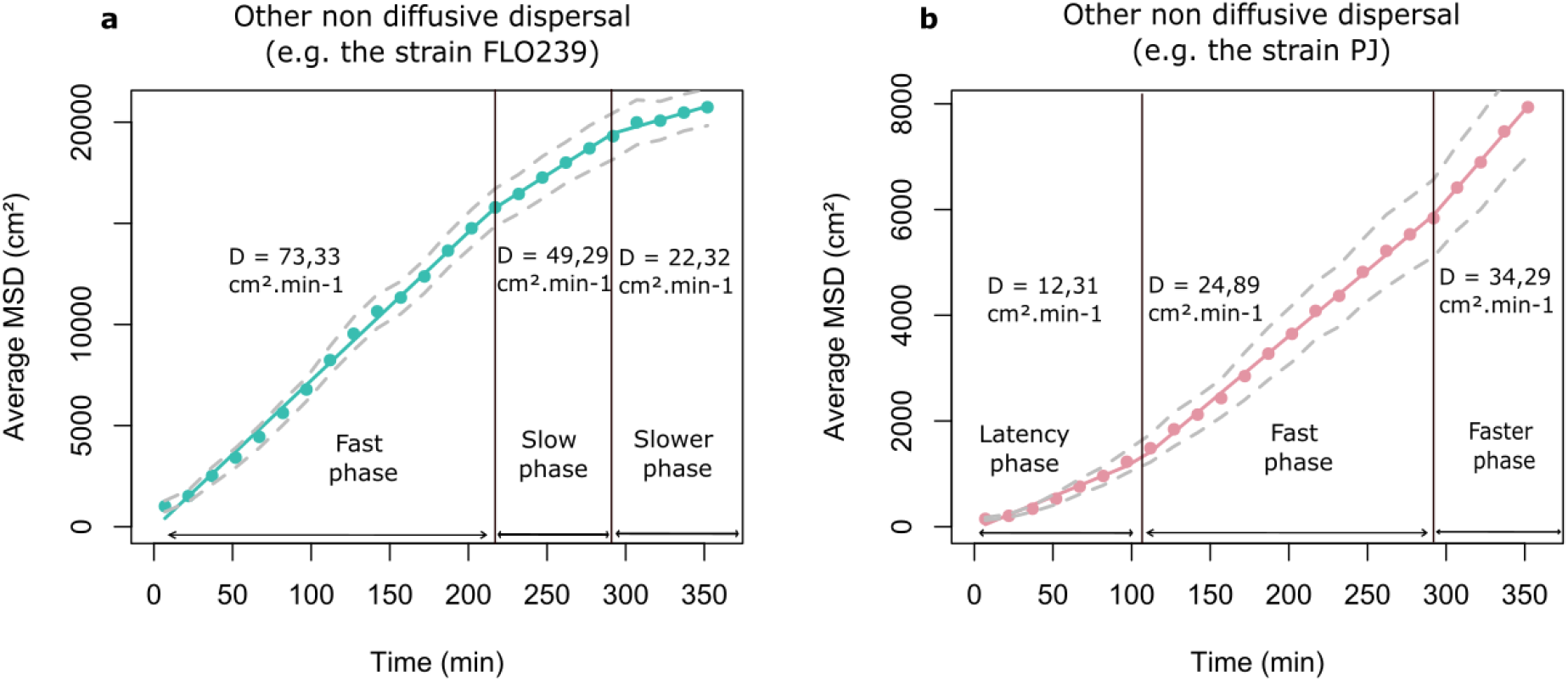
The two strains of the category “Others’’ in figure 7D which have a non-diffusive dispersal as with three distinct phases as in type 3, but specific patterns: FLO239 (a) is characterized by a first fast phase and then two successive slow phases. PJ (b), on the contrary, is characterized by a latency phase and two successive fast phases. Remark: we used piecewise linear regressions to test for deviations from linearity, but in those cases, it is possible that changes in diffusion coefficients are gradual rather than discrete: i.e. panel (a) might represent a gradual deceleration of spread, whereas panel (b) may feature a gradual acceleration of spread.

**Figure S3:**
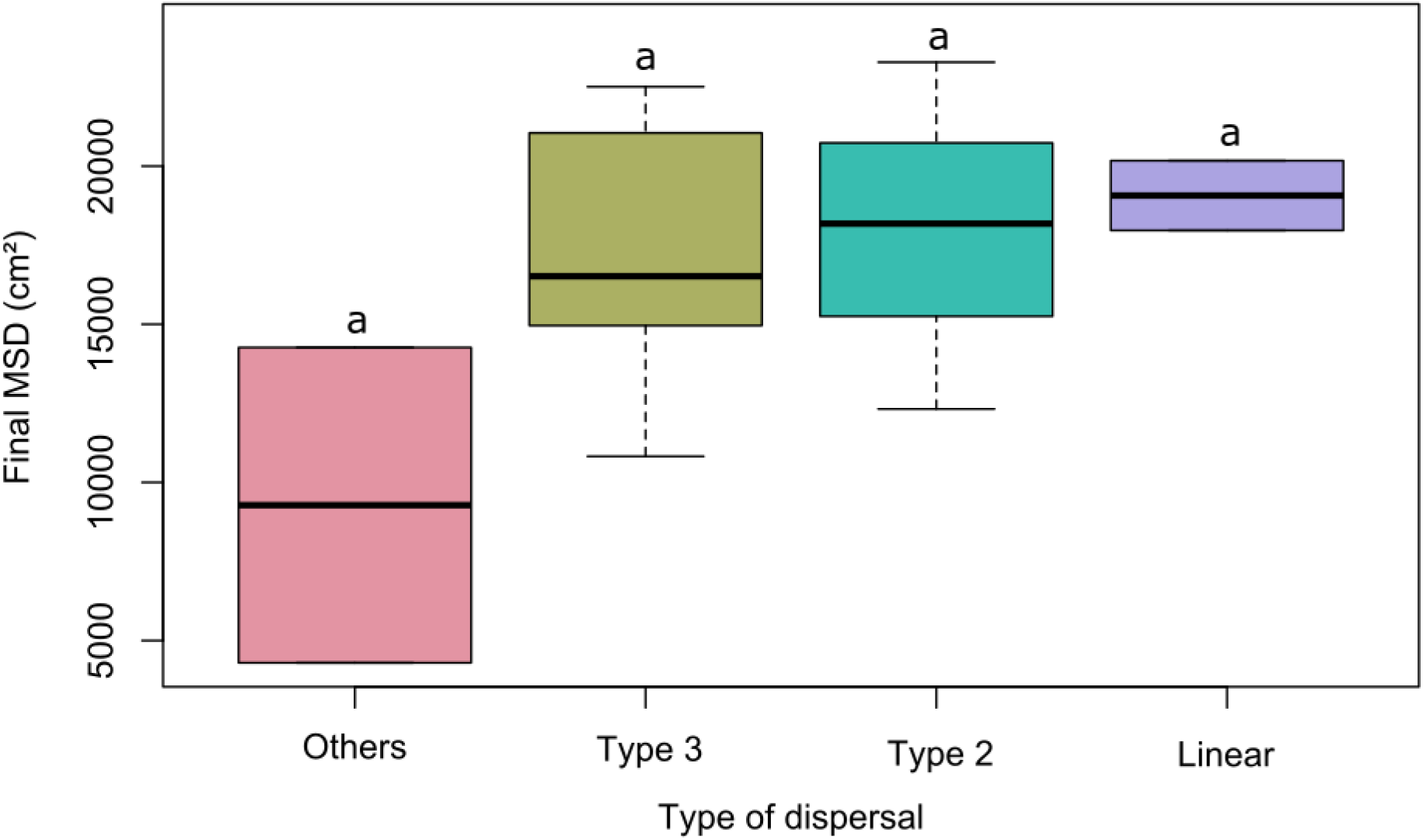
Boxplot of the final MSD according to the type of dispersal (anova; p = 0,19)

**Figure S4:**
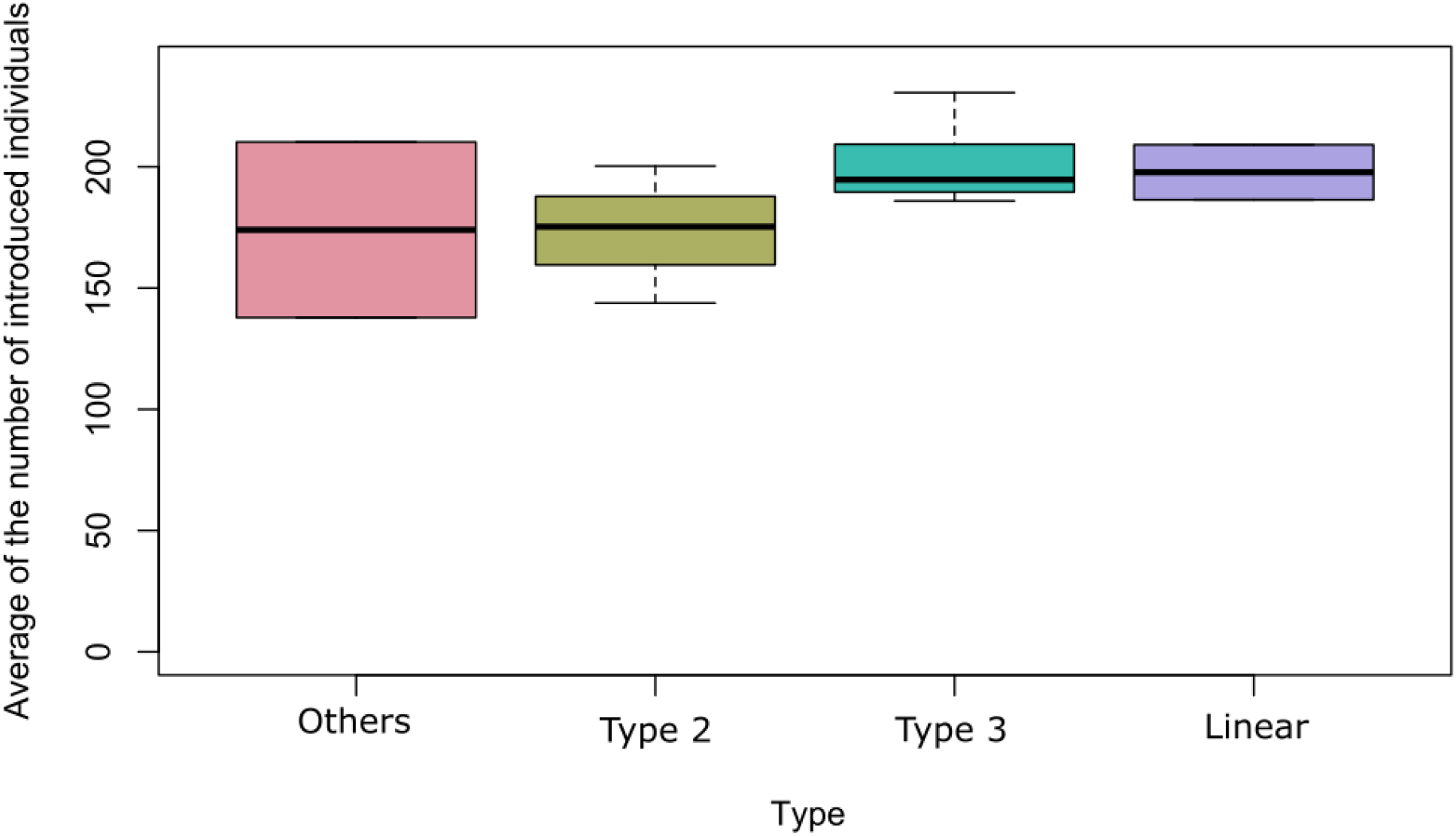
Boxplot of the average number of introduced individuals according to the type of dispersal. (anova; p=0.33)

